# Gene Gain and Loss from the Asian Corn Borer W Chromosome

**DOI:** 10.1101/2022.10.21.512844

**Authors:** Wenting Dai, Judith E. Mank, Liping Ban

**Author notes:** To whom correspondence should be addressed (LB).

## Abstract

We built a chromosome-level genome assembly of the Asian corn borer, *Ostrinia furnacalis* Guenée (Lepidoptera: Pyralidae, Pyraloidea), an economically important pest in corn, from a female, including both the Z and W chromosome. Despite deep conservation of the Z chromosome across Lepidoptera, our chromosome-level W assembly reveals little conservation with available W chromosome sequence in related species or with the Z chromosome, consistent with a non-canonical origin of the W chromosome. The W chromosome has accumulated significant repetitive elements and experienced rapid gene gain from the remainder of the genome, with most genes exhibiting pseudogenization after duplication to the W. The genes that retain significant expression are largely enriched for functions in DNA recombination, the nucleosome, chromatin and DNA binding, likely related to meiotic and mitotic processes within the female gonad.

## Introduction

The non-recombining, sex-specific portions of the genome, namely Y and W chromosomes, exhibit very different properties from the remainder of the genome^1^. Recent advances in sequencing have made it possible to assemble full, or nearly full, sequences of numerous Y chromosomes^2^, and these efforts have revealed general patterns such as the retention of genes related to dosage-sensitivity^3,4^ and male function^5-8^ as one might expect from a chromosome limited in its inheritance to males.

Studying Y chromosomes alone makes it difficult to differentiate the effects of sex-limitation from the effects of limitation to males, and so W chromosomes can be a powerful contrast to reveal the overall convergence of genomic patterns of these unusual regions of the genome. Despite providing an important comparison, sequencing of W chromosomes has lagged somewhat, with the most complete assemblies in birds^3,9^ suggesting that although W chromosomes share some similarities with Y chromosomes, namely an abundance of repetitive elements^10^, they may differ from Y chromosomes in that they lack genes with female-specific functions^3,9^. Whether this is unique to birds, or is a broader pattern of W chromosomes, remains unclear.

Lepidoptera, butterflies and moths, provide a key additional female heterogametic system^11,12^. The conservation of the Z chromosome has been well established in Lepidoptera^13^, however, the W chromosome in Lepidoptera is unusual in that it was recruited into the genome well after the origin of the Z chromosome^14^, as the basal lineages in the clade are Z0/ZZ. Available evidence suggests that, at least in some lineages, the W chromosome bears no homology to the Z^13,15,16^, and may actually have originated from a B chromosome^13^. Within the Lepidoptera, at the base of the Ditrysia group, complex sex chromosomes, including neo-W chromosomes are observed in many clades based on cytogenetic analysis^14^.

Recently, third-generation sequencing advances have permitted partial or chromosome-level assemblies of the W chromosome in a limited number of Lepidoptera, including *Cydia pomonella* (Torticidae, Tortricoideae)^16^, *Cnaphalocrocis medinalis* (Crambidae, Pyraloidea)^17^, *Spodoptera exigua* (Noctuidea, Noctuoidea)^18^, *Spodoptera frugiperda* (Noctuidea, Noctuoidea)^19^, *Trichoplusia ni* (Noctuidea, Noctuoidea)^20^ and *Dryas iulia* (Nymphalidae, Papilionoidea)^15^. This work has revealed no detectible homology between the Z and W in *C. pomonella*^16^, *T. ni*^21^ and *D. iulia*^15^. In contrast, substantial synteny is observed between the W chromosomes of *S. exigua* and *C. medinalis*^17,22^. All this is consistent with a non-canonical origin of the W chromosome in Lepidoptera, where the W has been recruited from a B chromosome and is therefore not homologous to the deeply conserved corresponding Z chromosome.

To answer questions about the conservation and gene content of the lepidopteran W, we built a chromosome-level genome of the Asian corn borer *Ostrinia furnacalis* Guenée (Lepidoptera: Pyralidae, Pyraloidea), is a major insect pest of corn, widespread in the Asian-Western Pacific region, using a combination of PacBio HiFi (CCS) sequencing and Hi-C. We couple this with extensive RNA-Seq analysis of multiple developmental stages and tissues in both sexes. Our catalog of W gene content reveals extensive duplications from all other chromosomes in the genome, with a relatively low percent of genes with persistent gene activity, which are enriched for functions in DNA recombination, the nucleosome, chromatin and DNA binding. Our results suggest that the W chromosome retains gene related to meiotic and mitotic functions within the female gonad, in contrast to the avian W chromosome^3,9^ and more consistent with findings from Y chromosomes^5-8^.

## Methods

### Samples

*O. furnacalis* larvae were collected from a corn field at Beijing, China, in July 2020, and fed with an artificial diet in the laboratory of China Agricultural University. The incubator environment was set at 26 ± 1℃ and 50 ± 5% relative humidity on a photoperiod (Light: Dark = 16:8).

### Genome sequencing and assembly

Genomic DNA was extracted from a female laboratory-fed pupa. From this single individual, we obtained 32.96 Gb circular consensus sequencing (CCS) data from one cell of the PacBio Sequel II platform at Biomarker Co., Ltd., Qingdao, China (Table S1). High accuracy PacBio Hifi data were assembled using hifiasm (v0.12) software to obtain a draft genome^23^. We used purge_haplotigs^24^ to remove redundant contigs and generate a non-redundant assembled genome.

### Hi-C scaffolding

We constructed Hi-C libraries (300-700 bp insert size), following Rao et al.^25^, and sequenced them with pair-end 150 Illumina Hiseq at Biomarker Co., Ltd., Qingdao, China. 53.05 Gb of clean data were produced after filtering adapter sequences, primer sequences and low-quality data (Table S2). The resulting trimmed reads were aligned to the draft assembly with BWA (v0.7.10-r789), retaining only uniquely aligned read pairs with mapping quality >20 for further analysis. Invalid read pairs, including dangling-end, self-cycle, re-ligation and dumped products, were removed by HiC-Pro (v2.8.1)^26^.

The 56.25% of unique mapped read pairs represent valid interaction pairs and were used for correction, clustering, ordering and orientation of scaffolds into chromosomes with LACHESIS^27^. Before chromosome assembly, we performed a preassembly for error correction of contigs which required the splitting of contigs into segments of 50 kb on average. Then Hi-C data were mapped to these segments using BWA (v0.7.10-r789). The uniquely mapped data were retained for the assembly, and any two segments which showed inconsistent connection with information from the raw scaffolds were checked manually. Parameters for running LACHESIS included: CLUSTER_MIN_RE_SITES = 27; CLUSTER_MAX_LINK_DENSITY = 2; ORDER_MIN_N_RES_IN_TRUNK = 15; ORDER_MIN_N_RES_IN_SHREDS = 15. After this step, placement and orientation errors exhibiting obvious discrete chromatin interaction patterns were manually adjusted. Finally, we constructed a heatmap based on the interaction signals of valid mapped read pairs between chromosomes.

### Genome annotation

Transposable elements (TEs) were identified by a combination of homology-based and de novo approaches. We first carried out a de novo repeat prediction using RepeatModeler2 (v2.0.1)^28^, which implements RECON (v1.0.8)^29^ and RepeatScout (v1.0.6)^30^. Then the predicted results were classified using RepeatClassifier^28^ by means of repbase (v19.06)^31^, REXdb (v3.0)^32^ and Dfam (v3.2)^33^. Secondly, we performed a de novo prediction for long terminal repeats (LTRs) using LTR_retriever (v2.8)^34^ via LTRharvest (v1.5.9)^35^ and LTR_finder (v2.8)^36^. A non-redundant species-specific TE library was constructed by combining the de novo TE library above with the known databases. Final TE sequences in the *O. furnacalis* genome were identified and classified by homology search against the library using RepeatMasker (v4.10)^37^. Tandem repeats were annotated by Tandem Repeats Finder (TRF v409)^38^ and MIcroSAtellite identification tool (MISA v2.1)^39^.

We integrated de novo prediction, homology search, and transcript-based assembly to annotate protein-coding genes. The de novo gene models were predicted using Augustus (v2.4)^40^ and SNAP (2006-07-28)^41^, both ab initio gene-prediction approaches. For the homolog-based approach, we used GeMoMa (v1.7)^42^ with reference gene models from *Drosophila melanogaster, Bombyx mori, Chilo suppressalis* and *Galleria mellonella*. For the transcript-based prediction, RNA from a mixture sample containing eggs, larvae, pupae and adults whole body of females and males was used for generating sequencing libraries and was sequenced on an Illumina NovaSeq 6000 platform (25.82-fold coverage of the genome, Table S3). RNA-sequencing data were mapped to the reference genome using Hisat (v2.0.4)^43^ and assembled with Stringtie (v1.2.3)^44^. GeneMarkS-T (v5.1)^45^ was used to predict genes based on the assembled transcripts. PASA (v2.0.2)^46^ was used to predict genes based on the unigenes assembled by Trinity (v2.11)^47^. Gene models from these different approaches were combined with EVM (v1.1.1)^48^ and updated by PASA (v2.0.2)^46^. The final gene models were annotated by searching the GenBank Non-Redundant (NR, 20200921), TrEMBL (202005), Pfam (v33.1)^49^, SwissProt (202005)^50^, eukaryotic orthologous groups (KOG, 20110125), gene ontology (GO, 20200615) and Kyoto Encyclopedia of Genes and Genomes (KEGG, 20191220) databases.

We used tRNAscan-SE (v1.3.1)^51^ with default parameters to identify transfer RNAs (tRNAs), and barrnap (v0.9) with default parameters to identify the ribosomal RNAs (rRNAs) based on Rfam (v12.0)^52^. miRNAs were identified with miRbase^53^. Small nucleolar RNA (snoRNAs) and small nuclear RNA (snRNAs) were identified with Infernal (v1.1.1)^54^, using the Rfam (v12.0) database^52^.

Pseudogenes have similar sequences to functional genes, but may have lost their biological function because of mutations. GenBlastA (v1.0.4)^55^ was used to scan the whole genome after masking predicted functional genes. Putative candidates were then analyzed by searching for premature stop codon and frameshift mutations using GeneWise (v2.4.1)^56^.

### Comparative genomics and phylogenetic reconstruction

Protein sequence alignments between *O. furnacalis* and seven other lepidopteran species (*B. mori, C. suppressalis, C. pomonella, C. medinalis, S. exigua, Spodoptera fugiperda,* and *T. ni*) were performed with diamond (-e < 1e-5), then alignment results were analyzed and the homologous chromosomal regions were identified with MCScanX (MATCH_SCORE: 50, MATCH_SIZE: 5, GAP_PENALTY: −1, OVERLAP_WINDOW: 5, E_VALUE: 1e-05, MAX GAPS: 15). The synteny relationships among chromosomes were displayed using circos (v0.69–9).

We used the protein sequences of *O. furnacalis and* ten other insect species (*S. frugiperda, Spodoptera litura, B. mori, C. suppressalis, Danaus plexippus, Melitaea cinxia, Papilio Xuthus, C. pomonella, Plutella xylostella* and *D. melanogaster*) for phylogenetic analysis, keeping only the longest transcript of each gene for analysis, and using OrthoFinder (v2.5.4)^57^ with default settings to identify orthologues and homologues. To infer the phylogeny of these insects, multiple sequence alignments of single-copy orthologs were performed using MAFFT (v7.471)^58^. Then we extracted conserved sequences with gblocks (v0.91b)^59^ and concatenated them to a single sequence alignment. The resulting sequence alignment was used to construct a maximum likelihood phylogenetic tree using IQ-TREE (v1.6.12)^60^ (outgroup: *D. melanogaster*). Divergence times between various species were estimated by MCMCtree in PAML (v4.9j)^61^. Three standard divergence time points from the TimeTree database (http://timetree.org/) were used for calibration: *S. frugiperda* – S. *litura*, 100-122 million years ago (Mya), *D. plexippus* – *M. cinxia*, 76-102 Mya, (*D. plexippus, M. cinxia*) – *P. xuthus*, 76-146 Mya. The tree was visualized using FigTree (v1.4) (http://tree.bio.ed.ac.uk/software/figtree/). The gene count table from OrthoFinder (v2.5.4) was used as inputs to examine the expansion and contraction of each gene family in cafe (v5.1.0)^62^.

### Detection of sex chromosomes

In order to identify the sex chromosomes, we performed genome resequencing with five female and male pupae. High-quality clean data (24.95-28.80-fold coverage of the genome) were obtained through the pair-end 150 Illumina Hiseq platform at Biomarker Co., Ltd., Qingdao, China (Table S3). The clean data were aligned to reference genome with Bwa-men (v0.7.17). We compared the coverage differences between male and female samples^63^ to distinguish the Z, W and autosomes. Specifically, we used the genomecov and groupby in Bedtools (v2.30.0) to obtain a per-base median coverage depth for each chromosome and normalized them by the mean of all chromosome median coverages for each sample. Normalized coverage depth was averaged by sex to produce a coverage depth per chromosome for each sex. Then we compared coverage depth between sexes for each chromosome and calculated log_2_ male:female (M:F) coverage ratio [log_2_(M:F coverage)]. Autosomes have an equal coverage between sexes [log_2_ (M:F coverage) = 0], while the Z chromosome should show approximately two-fold greater coverage in the males [log_2_(M:F coverage) = 1]. The W chromosome should show a strong female-biased coverage pattern. We also calculated the M:F median coverage ratio along each chromosome with nonoverlapping 1000 bp windows.

### W-gene homologs search and calculation of synonymous divergence (d*_S_)*

Predicted protein sequences were used to detect reciprocal best hits between the W chromosome and the remainder of the genome using getRBH.pl (https://github.com/Computational-conSequences/SequenceTools) for *O. furnacalis*. For each pair of orthologous genes, we deleted stop codes and aligned the coding sequences using macse (v2.05)^64^, then extracted conserved sequences with gblocks (v0.91b)^59^. The resulting alignments were used as inputs of CODEML in PAML (v4.9j)^61^ to estimate pairwise synonymous divergence with settings runmode = −2, seqtype = 1 and CodonFreq = 2. Since divergence estimates are not reliable for saturated sites, we excluded orthologs with *d_S_* > 3^65^.

The homolog search of *O. furnacalis* W in *C. pomonella, C. medinalis, S. exigua, S. fugiperda* and *T.ni* genome also used getRBH.pl. We assessed enrichment of Gene Ontology^66,67^ terms for W gene content.

### W-gene expression level

We used two different RNA-Seq datasets to determine W expression, the mixture sample of RNA-Seq data from eggs, larvae, pupae and adults used for genome annotation (described above), and data from female gonad, as sex-limited chromosomes are often largely expressed in the gonad. For female gonad, RNA from five gonad samples of adult females was sequenced on an Illumina NovaSeq 6000 platform (10.14-fold coverage of the genome, Table S3). For each dataset separately, we used fastp (v0.20.0)^68^ to filter out low quality reads and to remove adapters with the default parameters. Then we mapped the clean reads to the *O. furnacalis* reference genome using HISAT2 (v2.1.0)^43^. The FPKM (fragments per kilobase of exon per million fragments mapped) of each gene was determined using Stringtie (v2.1.4)^44^ based on the annotated GFF file. For female gonad, we used averaged FPKM of five samples.

### Copy number variation of W and Z/autosome genes

The amino acid sequences of protein-coding genes from whole genome were used as the input of the blastp mapping against the Swiss-Prot database to assign gene symbols (abbreviations for the gene names). We BLASTed each W gene to the remainder of the genome to identify the best hit and only retained orthologous genes on the W chromosome and Z/autosomes that consistently mapped to the same protein (reciprocal best BLAST hit) and calculated their copy number following the protocol in Mueller et al.^69^.

## Results

### Chromosome-level genome assembly of the Asian Corn Borer

Our chromosome assembly strategy employed PacBio HiFi (CCS) sequencing data to assemble the draft genome and Hi–C data to detect chromosomal contact information. These PacBio long-reads were self-corrected using Quiver and assembled into a draft genome assembly with a total length of 493.10Mb, consisting of 57 contigs with an N50 length of 15.69Mb (Table S4). The assembled genome size is similar to that obtained from genome surveys (Supplementary Figure 1). Next, Hi–C linking information further anchored, ordered, and oriented 46 contigs to 32 chromosomes (30 autosomes, with Z and W sex chromosomes), which contained 86.79% of the whole genome assembly (Table S4, Supplementary Figure 2). The chromosome-level genome assembly was 492.57Mb with a scaffold N50 of 16.47Mb (Table S4).

We evaluated the completeness of *O. furnacalis* genome assembly with the Metazoan Benchmark of Universal Single-Copy Orthologs (BUSCO v4) and CEGMA core genes (CEGMA v2.5) datasets. Our assembly contained 97.69% of BUSCO genes, of which 97.06% were single copy and 0.63% were duplicates, and 96.07% of CEGMA genes, of which 79.03% were highly conserved (Table S5). To further evaluate the genome assembly quality, genomic DNA sequencing data from Illumina HiSeq were mapped to the assembly scaffolds, with a 99.07% coverage rate (Table S5). Finally, lepidopteran genomes typically exhibit very high levels of synteny. Whole-genome alignment of the *O. furnacalis* assembly to the chromosomes of the *S. litura* revealed that chromosomal linkage and ordering of genes are highly conserved (Supplementary Figure 3). All these analyses indicated that the genome assembly is both highly reliable and complete for subsequent analyses.

### Genome annotation

In total, 860,391 repeat sequences spanning 204.4Mb were identified, constituting 41.45% of the *O. furnacalis* genome (Table S4, Figure 1a). We integrated the result of *ab initio*, homology-based and RNA-seq methods to annotate protein-coding genes, most of the which came from homology-based and RNA-seq methods, indicating the high-quality of annotation (Supplementary Figure 4). Finally, we obtained 16,509 protein coding genes in the *O. furnacalis* genome, which has similar gene features to other lepidopteran genomes (gene length, gene number, coding length, intron length of each gene, exon length) (Supplementary Figure 4). Next, we identified different types of noncoding RNAs (ncRNAs), including 7,710 transfer RNAs (tRNA), 73 ribosomal RNAs (rRNA), and 39 microRNAs (miRNA) (Table S6). In addition, we annotated 167 pseudogenes in the *O. furnacalis* genome, defined as any genomic sequence that is similar to another gene but is defective, such as containing a premature stop codon or a frameshifts mutation^70^. Most pseudogenes were located in the chromosome LG3 (Figure 1c, Table S6), the W chromosome (see below). We performed a functional annotation for all predicted protein coding genes using NCBI nonredundant, EggNOG, GO, KEGG, SWISS-PROT and Pfam databases, with 98.21% of our predicted genes assigned to at least one of these databases (Table S6).

**Fig 1.**
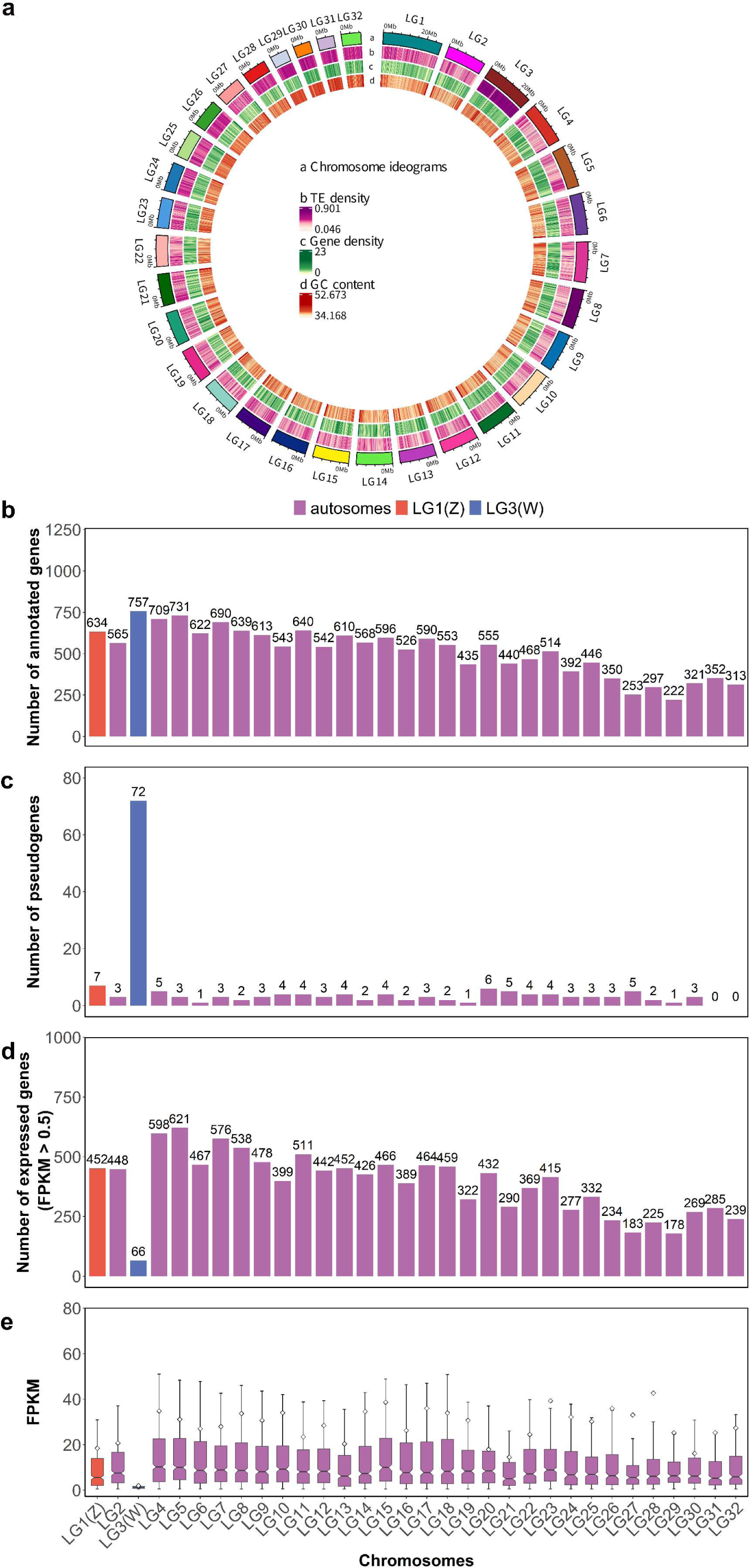
Genomic characterization of *Ostrinia furnacalis*. a. Circos plot depicting the genomic landscape of the 32 chromosomes (Chr1–Chr32 on a Mb scale). The denotation of each track is listed in the center of the circle. Blue lines in LG3 (the W chromosome) represent collinearity within the W, due to repeat elements. b. Number and distribution of protein-coding genes. c. Number and distribution of pseudogenes. d. Number of expressed genes (FPKM > 0.5 in mixed sample). e. Average chromosomal expression level (excluding genes with FPKM ≤ 0.5 in mixed sample).

We used OrthoFinder to find orthologous genes among *O. furnacalis* and ten other insect species (see Materials and Methods). A total of 16231 orthologous groups with 1163 single-copy orthologous genes were identified. We inferred a phylogenetic tree and divergence time estimate using the end-to-end concatenated amino acid sequences of 1163 single-copy orthologous genes. The Crambidae lineage to which *O. furnacalis* belongs was estimated to diverge from Obtectomeran Lepidoptera approximately 77.3 Mya ago (Figure 2a).

**Fig 2.**
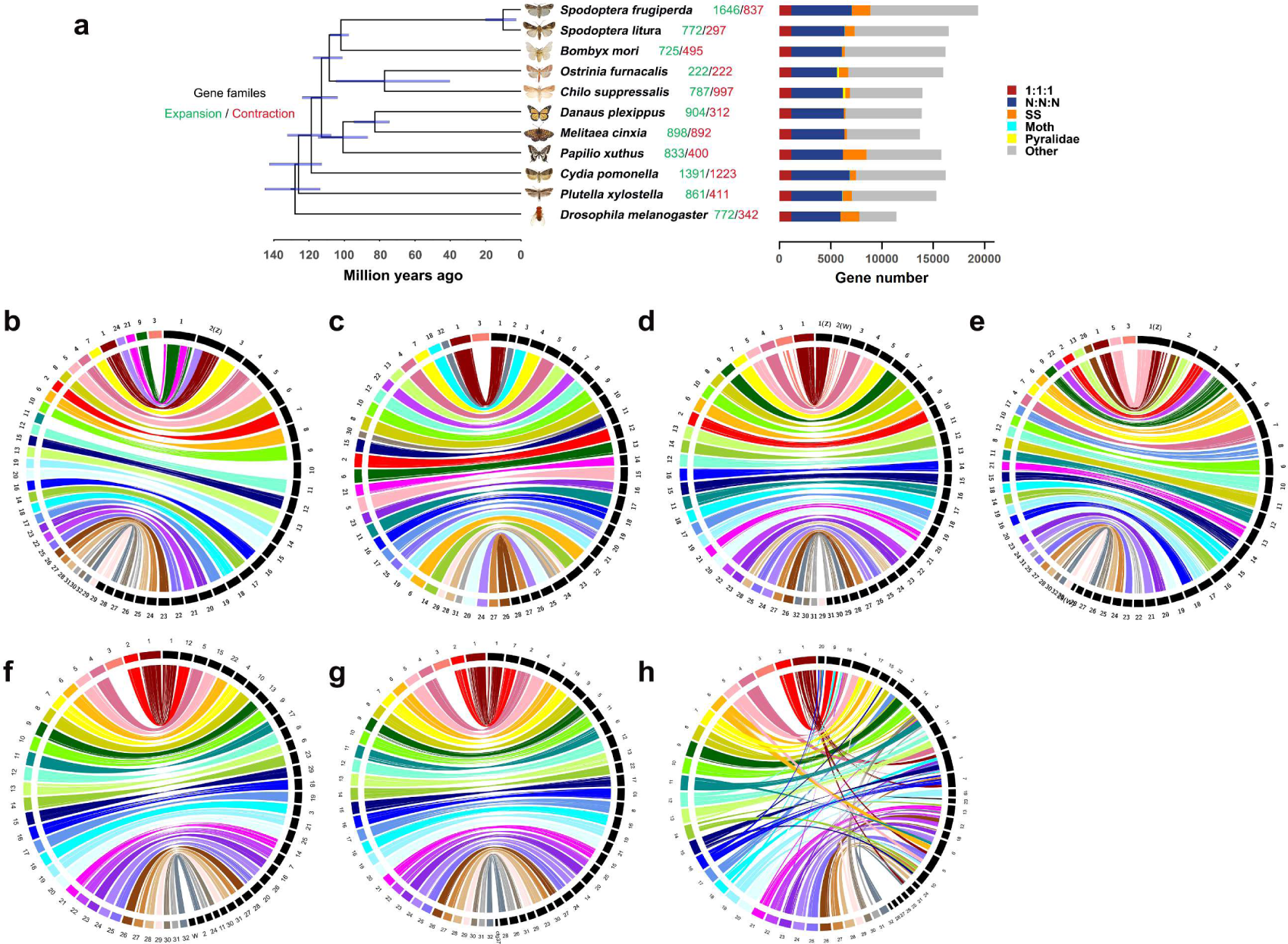
**a.** Phylogenetic tree and gene orthology of *O. furnacalis* with nine lepidopteran genomes and *D. melanogaster*. Comparative analysis of synteny between *O. furnacalis* and **b.** *C. suppressalis*, **c.** *B. mori*, **d.** *C. medinalis*, **e.** *C. pomonella,* **f.** *S. exigua,* **g.** *S. fugiperda*, and **h.** *T. ni*. The chromosomes of *O. furnacalis* are shown in the left, and the other insects’ chromosomes are shown in the right. The *O. furnacalis* W chromosome is LG3, and the Z is LG1 (b-h).

### Synteny, karyotype evolution, and sex chromosomes

We compared the syntenic relationships between *O. furnacalis* and other lepidopterans, including *B. mori, C. suppressalis, C. medinalis, S. litura, S. exigua, S. fugiperda, and T. ni* (Figure 2b-h). In general, the *O. furnacalis* genome shows a high level of synteny with other lepidopteran genomes, though with some fusion and fission events. *O. furnacalis* LG1 is syntenic with the Z chromosome in all other species, suggesting it is the Z chromosome in *O. furnicalis* as well, and this is also evident from M:F coverage analysis (Figure 3a), which reveals two-fold greater coverage in males (ZZ) compared to females (ZW).

**Fig 3.**
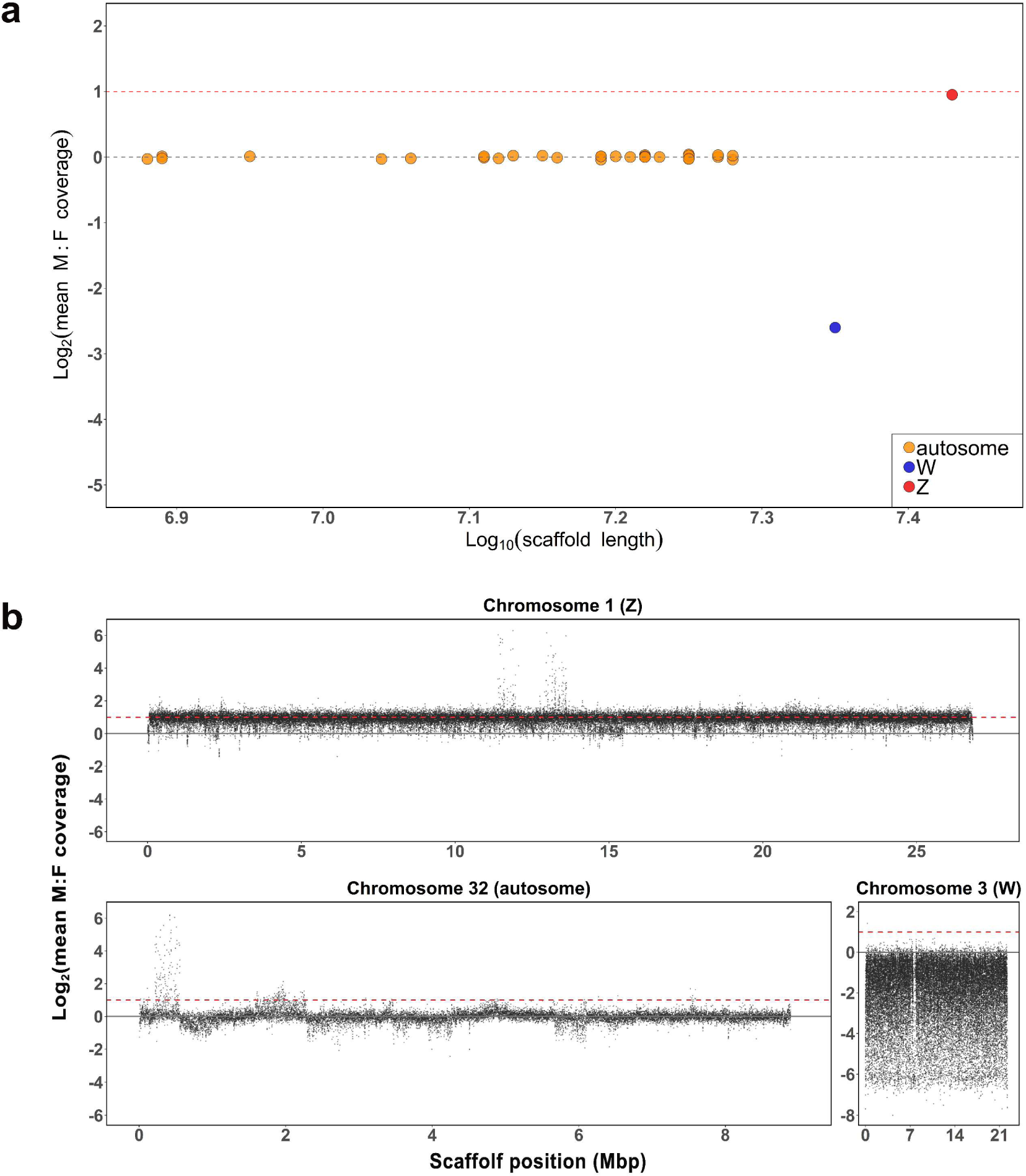
Detection of sex chromosomes. **a.** Male:female coverage ratios for each chromosome, plotted by chromosome length. Each point represents a single chromosome. The dashed gray line is the theoretical expectation for autosomes and the dashed red line shows the expectation for the Z chromosome. **b.** Male:female coverage ratios plotted in 500bp windows across scaffolds for Z, W, and a representative autosome (LG32).

*O. furnacalis* LG3, the W chromosome, exhibits female-biased coverage consistent with a female-limited chromosome (Figure 3a). The M:F coverage is highly variable across chromosome LG3 compared to all other chromosomes (Figure 3b), due in large part to the abundance of TEs on this chromosome (Supplementary Figure 5, Supplementary Figure 6). The W chromosome (LG3) is the second largest chromosome (22.23 Mb) in our assembly (Figure 1a), with the largest number of predicted protein coding genes (excluding pseudogenes) compared with other chromosomes (Figure 1b). The W also has the largest number of pseudogenes (Figure 1c), and contains 43.1% of all pseudogenes annotated in the genome. Many genes that are not technically pseudogenized were expressed below our 0.5 FPKM threshold. Only 66 protein coding genes showed significant expression above this threshold in our mixed sample (comprised of eggs, larvae, pupae and adults, see Materials and Methods, Figure 1d, 1e). 48 W genes showed expression >0.5 FPKM in adult female gonads.

Furthermore, LG3 has a notably greater repeat density and significantly different repeat composition compared to other chromosomes (Supplementary Figure 5, Supplementary Figure 6) with a larger proportion of satellites, DIRS, LINE, Copia and Gypsy, and an enrichment of *Maverick* elements.

### W homology

Given that previous reports have found no evidence of homology between the Z and W in some basal lepidopteran species^15,16,21^, we next investigated the evolutionary history of the *O. furnacalis* W chromosome. Using our W gene set, we first examined homology between the *O. furnacalis* W chromosome and the W chromosome from other lepidopterans. The *O. furnacalis* W shows substantial homology with the W in *C. medinalis*, consistent with some form of shared ancestry within the family Crambidae. However, we observed no discernable homology between the *O. furnacalis* W and the W chromosome of any of the more distantly related lepidopterans that we queried (Figure 2b-h).

Reciprocal best BLAST hits between the W coding catalog and the remainder of the *O. furnicalis* genome reveals similarity throughout the genome (Figure 4), consistent with a B chromosome origin of the W chromosome followed by gene duplication from locations throughout the genome. Of the 482 W coding genes having Z/autosome paralogs (Figure 4), 202 exhibited fewer or no introns in the W paralogs, consistent with retrotransposition.

**Fig. 4.**
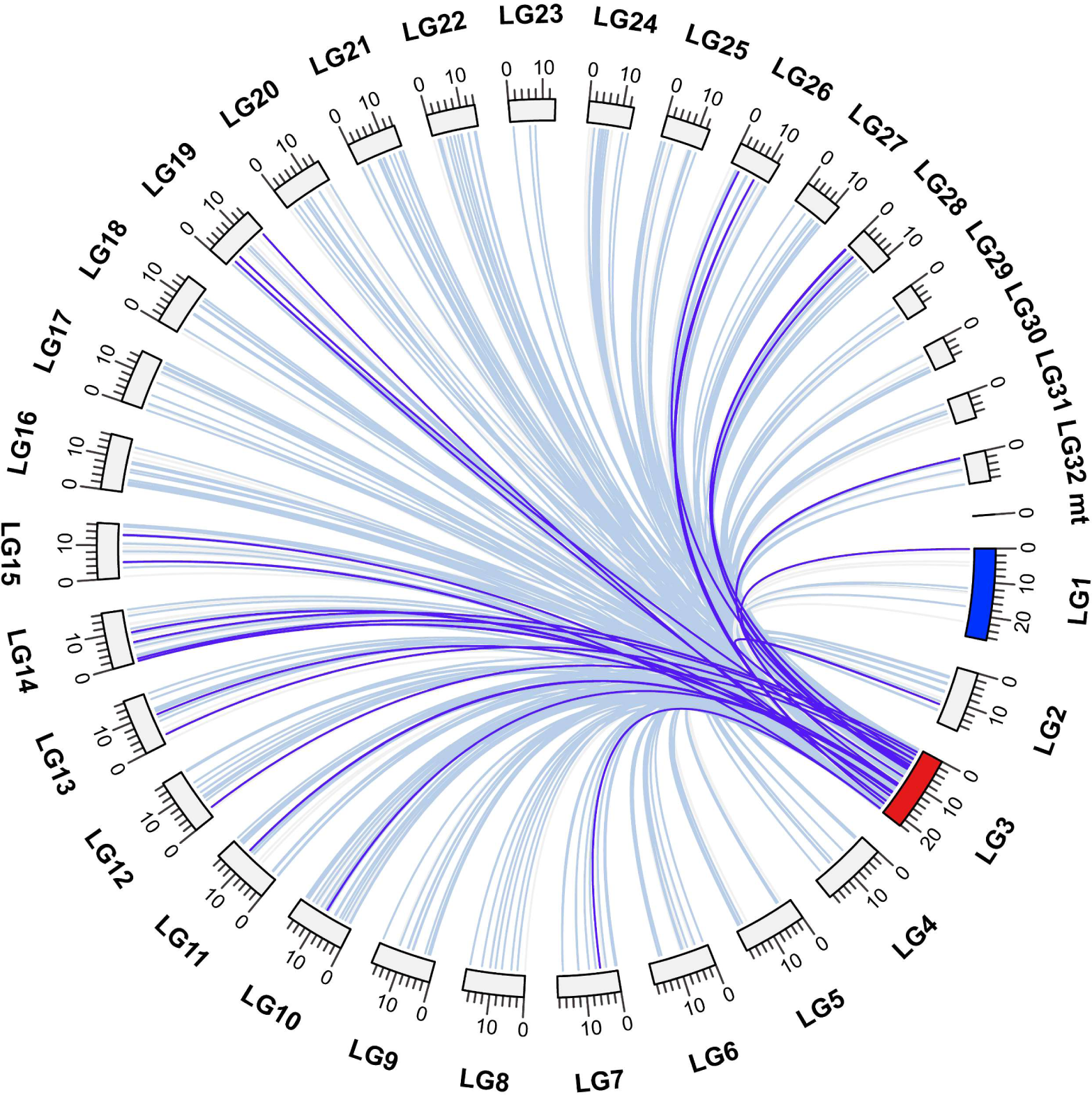
The reciprocal best hits search for W (LG3) genes. W chromosome (LG3) is marked in red and Z chromosome (LG1) in blue, mt represent the mitochondrial genome of *O. furnacalis.* Light blue line: identity ≥ 90%; purple line: 80% ≤ identity < 90%; light grey line: identity < 80%.

We next examined synonymous divergence (d_S_) for these paralogs in order to determine the timing of gene duplication to the W chromosome. The density of d_S_ (Figure 5a) shows that most paralogs with W genes have relatively low divergence, and the distribution of paralog d_S_ across the W chromosome suggests that duplications occur randomly throughout the chromosome. Given the low percent of genes on the W chromosome that exhibit expression >0.5 FPKM and the high number of pseudogenes on the W (Figure 1), it is likely that most duplicates to the W chromosome fail to maintain significant expression and are silenced relatively quickly.

**Fig 5.**
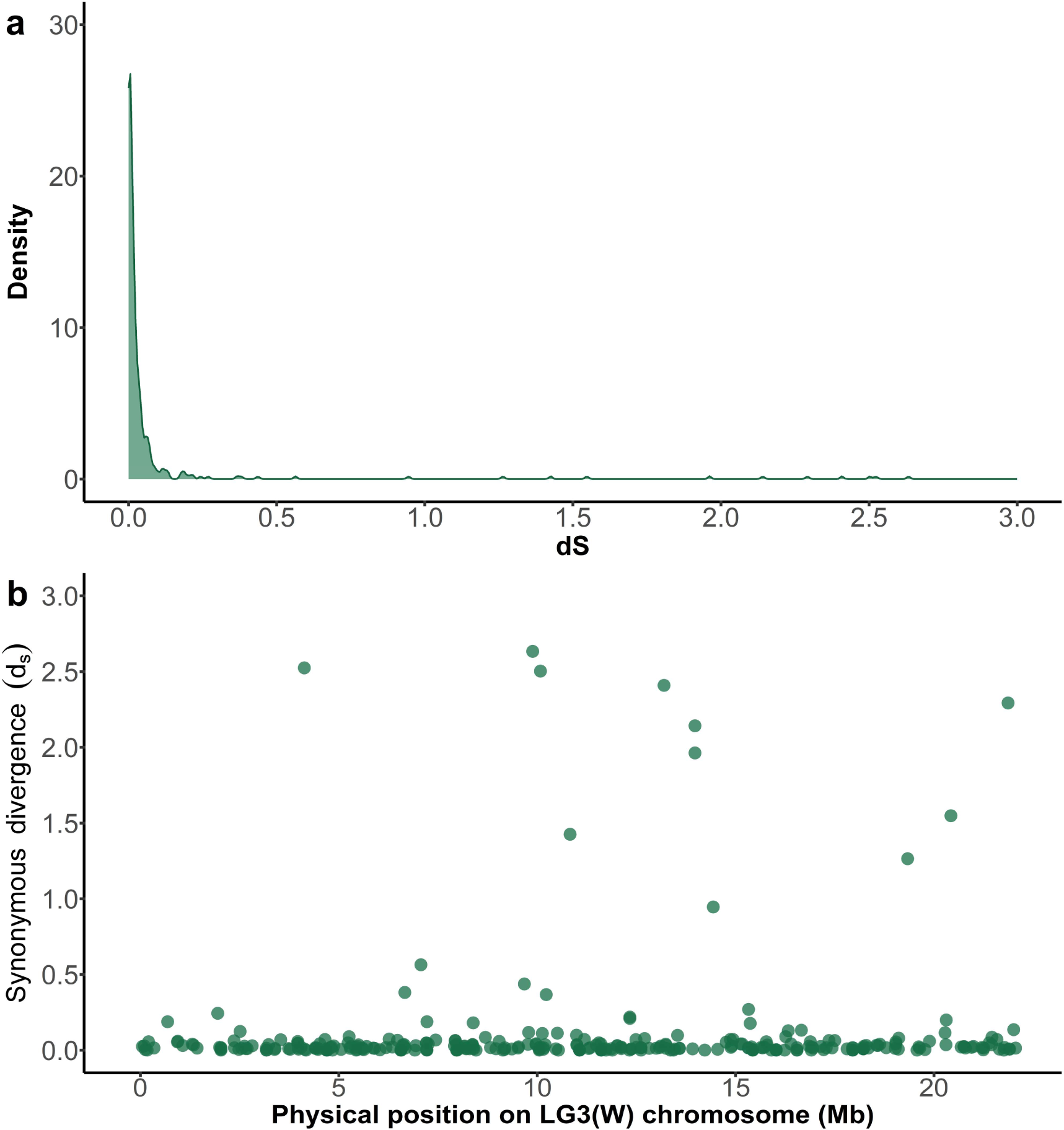
Synonymous divergence between W gene and best BLAST hit gene from the remainder of the genome. a. Density plot, b. Distribution across the W chromosome.

Sex-limited chromosomes often contain many copies of some genes^5,71^, and so we examined copy number for genes on the W chromosome and their most similar paralog (Figure 6, Table S7). Overall, we observed lower number of gene copies on the W than for the autosomal/Z chromosome paralog.

**Fig. 6.**
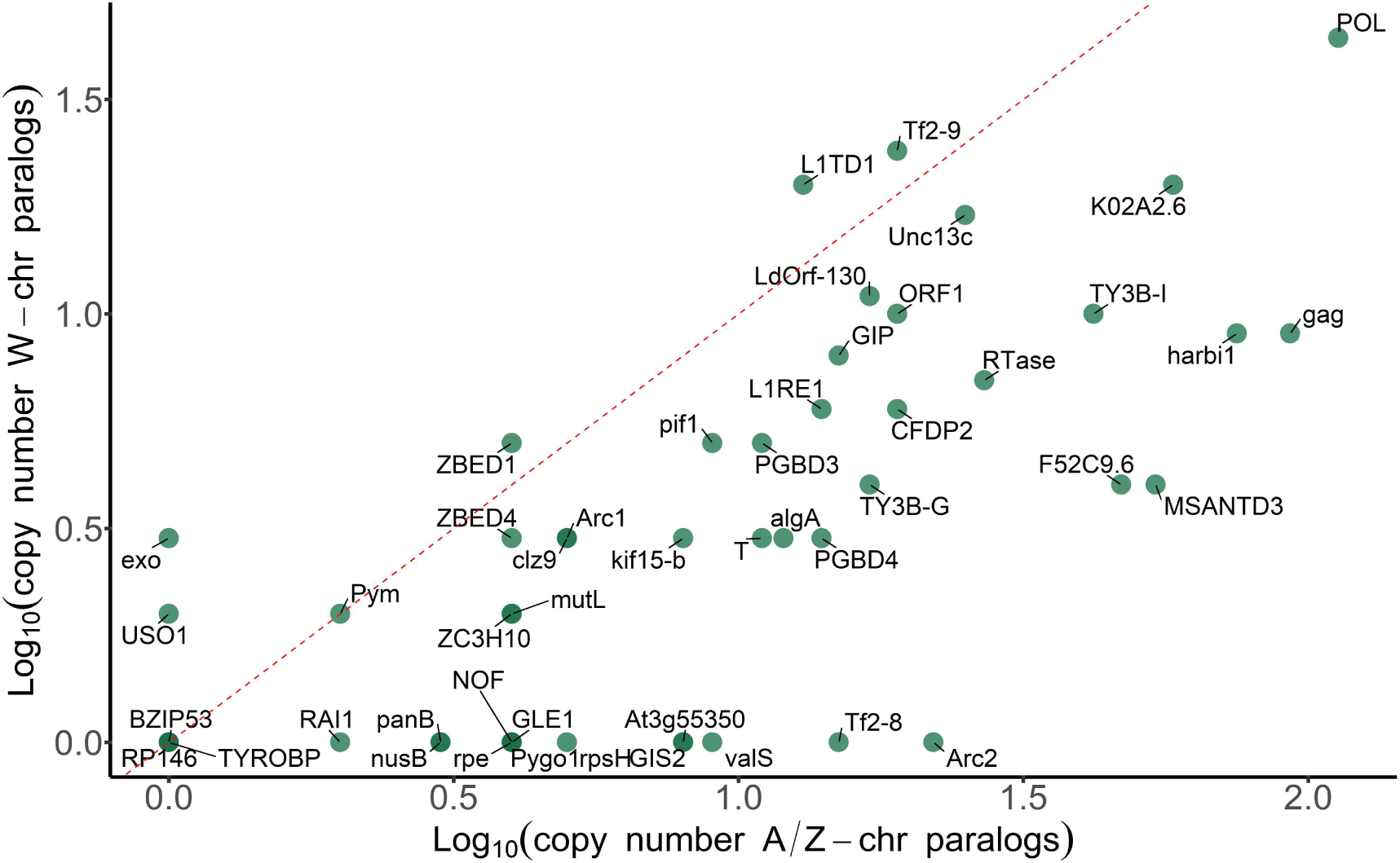
Copy number for W and autosomal/Z chromosome paralogs.

Given the rapid apparent decay of W chromosome paralogs (Figure 5), we examined the Gene Ontology enrichment for W expressed genes (>0.5 FPKM) (Tables 1 and 2), and observed statistical enrichment of terms related to functions in DNA recombination, the nucleosome, chromatin and DNA binding. Together, these results suggest that many genes retained with functional expression on the W chromosome are related to mitotic and meiotic processes, much of it within the female gonad.

**Table 1.** Functional Enrichment of Significantly Expressed Genes (FPKM > 0.5 from mixed sample) on *O. furnicalus* W chromosome

| PANTHER GO | FDR | Expressed W genes |
| --- | --- | --- |
| **Biological** |  |  |
| Negative regulation of DNA recombination | 7.64 e-11 | 5 |
| Nucleosome assembly | 8.35 e -10 | 5 |
| Chromosome condensation | 2.48 e-8 | 5 |
| Telomere maintenance | 2.67 e-02 | 2 |
| **Molecular** |  |  |
| Nucleosomal DNA binding | 1.16 e-11 | 5 |
| Structural constituent of chromatin | 1.13 e-10 | 5 |
| Double-stranded DNA binding | 1.66 e-7 | 7 |
| **Cellular** |  |  |
| Nucleosome | 1.68 e-10 | 5 |
| Chromatin | 6.12 e-7 | 6 |

**Table 2.** Functional Enrichment of Significantly Expressed Genes (FPKM < 0.5 from female gonad) on *O. furnicalus* W chromosome

| PANTHER GO Term | FDR | Expressed W genes |
| --- | --- | --- |
| **Biological** |  |  |
| Negative regulation of DNA recombination | 2.04e-10 | 5 |
| Nucleosome assembly | 2.22e-9 | 2 |
| Chromosome condensation | 7.70e-08 | 5 |
| **Molecular** |  |  |
| Nucleosomal DNA binding | 3.09e-11 | 5 |
| Structural constituent of chromatin | 3.02e-10 | 5 |
| Double-stranded DNA binding | 8.28e-05 | 6 |
| **Cellular** |  |  |
| Nucleosome | 4.46e-10 | 5 |
| Chromatin | 3.19e-6 | 6 |

## Discussion

Here we report a high quality, chromosome-level genome for *O. furnicalis*. Our reference genome includes a single, contiguous sequence for the female-specific W chromosome, allowing us to query the content of this unique region of the genome.

W and Y chromosomes are often enriched for pseudogenes, as the lack of recombination in these regions leads to high rates of gene silencing^1^. At the same time, these regions often accumulate repeat sequences^9,72^, resulting in significantly higher repeat content compared to other chromosomes^9,16,17^. Consistent with this, we although we found that the number of annotated protein-coding genes of W chromosome is indeed the largest of all the chromosomes in the genome (Figure 1), most of these are pseudogenes (Figure 1c), and those genes that retain functional coding sequence have low overall transcriptional activity (Figure 1d)^12^.

We also found the W is enriched for repetitive elements in *O. furnicalis* (Supplementary Figure 5 and Supplementary Figure 6), with the number of *Maverick* elements particularly higher on the W chromosome compared to the rest of the genome (Supplementary Figure 5). *Maverick*, also known as *Polinton*, is a DNA transposon widespread in eukaryotes^73^, and one of the most striking characteristics of *Maverick* is the large size, typically 15–20 kb^73^.

### Conservation of sex chromosomes across Lepidoptera species

We searched orthologs to Z-linked genes of *O. furnacalis* in other lepidopterans, and found that although the Z chromosome shows clear strong conservation^14^ (Figure 2) the *O. furnacalis* W chromosome is only conserved with *C. medinalis*, also a member of Crambidae (Figure 2b-h). This is consistent with previous work suggesting that the composition of the W varies even among species with the same family^74^.

### Evolutionary history of the sex chromosomes in *O. furnacalis*

Karyotype work has revealed a complex history of the W chromosome, including the repeated origin of neo-W chromosomes in many lepidopteran lineages. The evolutionary history of the W chromosome in Lepidoptera differs from the canonical model of sex chromosome formation in that it was recruited, possibly from a B element, after the formation of the Z chromosome^14^, at the common root of Ditrysia and Tischerioidea^12,75^. Lineages ancestral to this recruitment exhibit Z0:ZZ karyotypes^14^. The B-chromosome origin of the W is supported by the fact that in some basal lineages, W chromosome bears no homology to the Z^13,15,16^.

We observe little similarity in gene content between the W and Z chromosome (Figure 4), a steady rate of gene duplication to the W chromosome from throughout the genome (Figure 5), and reduced intron number for many W paralogs compared to their most similar autosomal/Z-linked copy suggests that this likely often occurs through retrotransposition across the W chromosome. Genes retrotransposed or otherwise duplicated to the W chromosome will immediately experience the full degenerative effects of a non-recombining region^1,76^, and consistent with this, we observe high levels of pseudogenization (Figure 1) on the W chromosome. However, those genes on the W that retain functional coding sequence and expression (e.g. non-pseudogenized) are enriched for mitotic and meiotic functions, and display Gene Ontology term enrichments related to recombination, chromosome packaging and replication (Tables 1 and 2). Although it has been suggested that W chromosomes may differ from Y chromosomes in that they do not acquire genes related to female gonadal function^3,9^ in the same way that Y chromosomes retain genes related to male function^5-8^, our results suggest that this may not be a generalized pattern of W chromosomes. Indeed, the W chromosome in *O. furnicalis* are enriched for functions related to female meiosis and mitosis.

## Authors’ contributions

LB conceived and designed the research. WD performed the experiments, analyzed and interpreted the data, and wrote the manuscript. JEM helped analyze the data and write the manuscript. All authors read and approved the final manuscript.

## Acknowledgments

This work was supported by the Beijing Agriculture Innovation Consortium (BAIC02-2023) and National Key R&D Program of China (2022YFD1400500). JEM gratefully acknowledges support from NSERC and a Canada 150 Research Chair. This work was supported by the high-performance computing platform of China Agricultural University.

