## Supplementary Tables and Figures for "Gene Gain and Loss from the Asian Corn Borer W Chromosome"

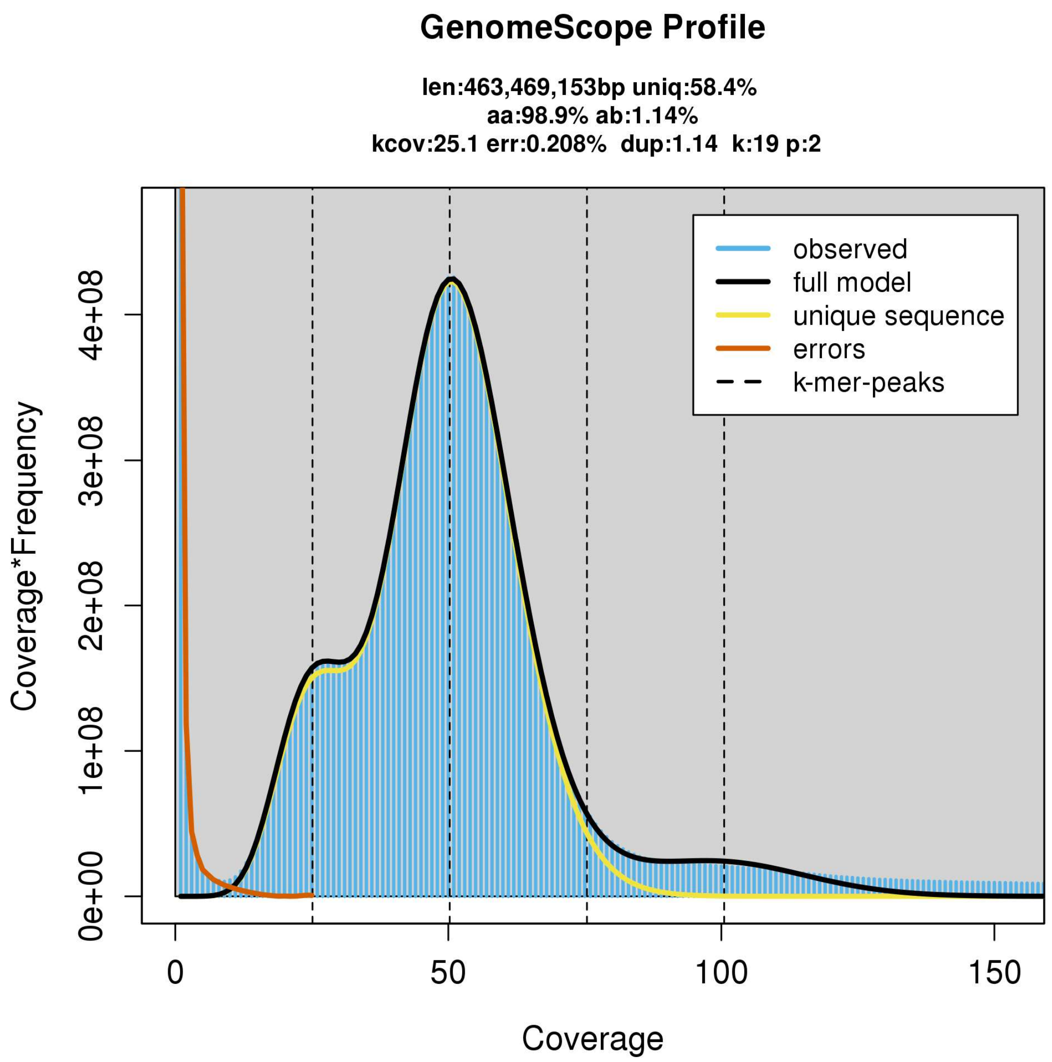


**Supplementary Figure 1. Genome survey of *Ostrinia furnacalis* using k-mer analysis.**

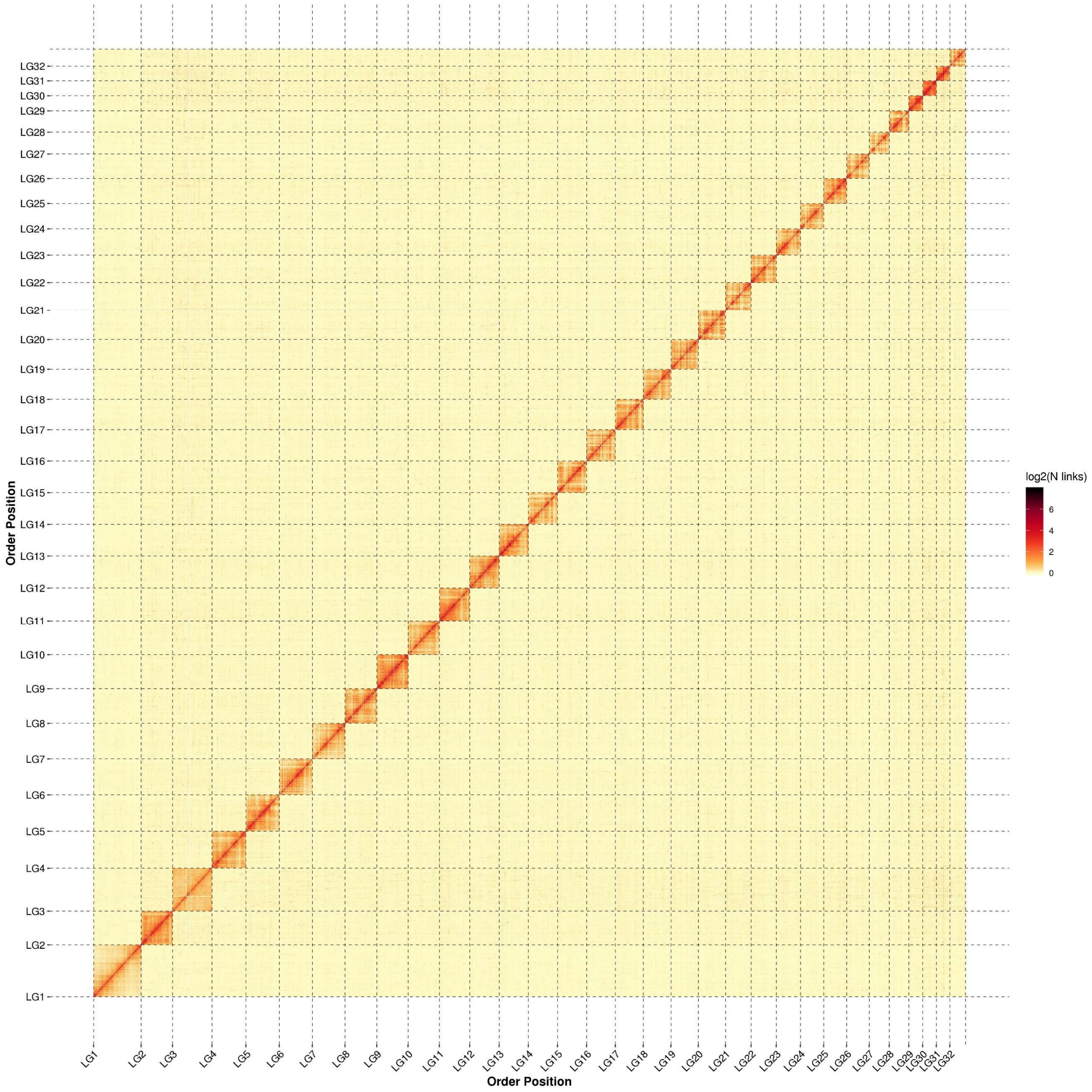


**Supplementary Figure 2. The genome-wide Hi-C interaction maps of 32 chromosomes in *Ostrinia furnacalis*.** The map indicates that intrachromosome interactions were strong while interchromosome interactions were weak. The shading gradient represents the chromosome interactions.

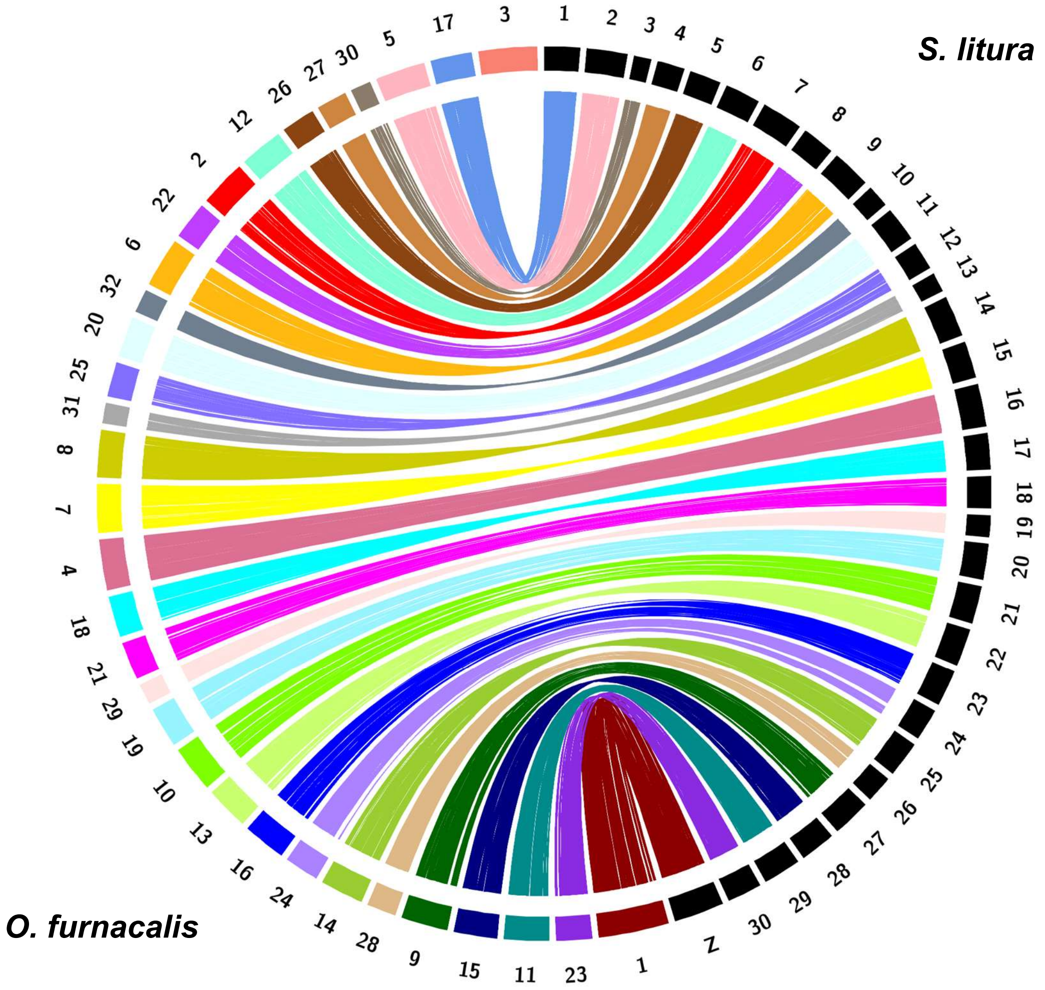


**Supplementary Figure 3. Synteny analysis of between *Ostrinia furnacalis* and *Spodoptera litura* chromosomes. Chromosomes of *Ostrinia furnacalis* are shown in the left, number 3 represent the W chr and 1 represents the Z chr. The chromosomes of *Spodoptera litura* are shown in the right.**

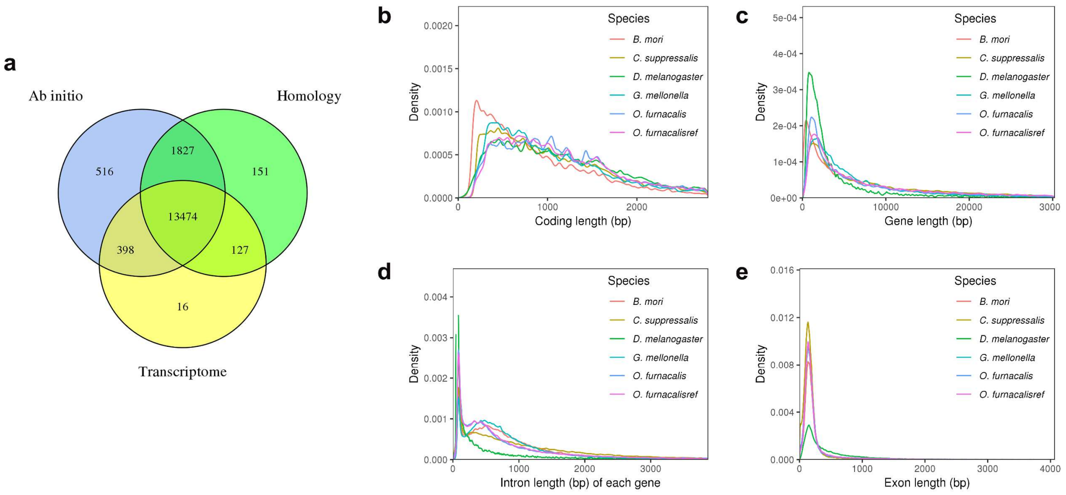


**Supplementary Figure 4. Annotation and evaluation of protein-coding genes. a.** Genes annotated via Ab initio, Homology-based and RNA-seq methods. **b-e.** Comparison of *Ostrinia furnacalis* gene features with other lepidopteran genomes.

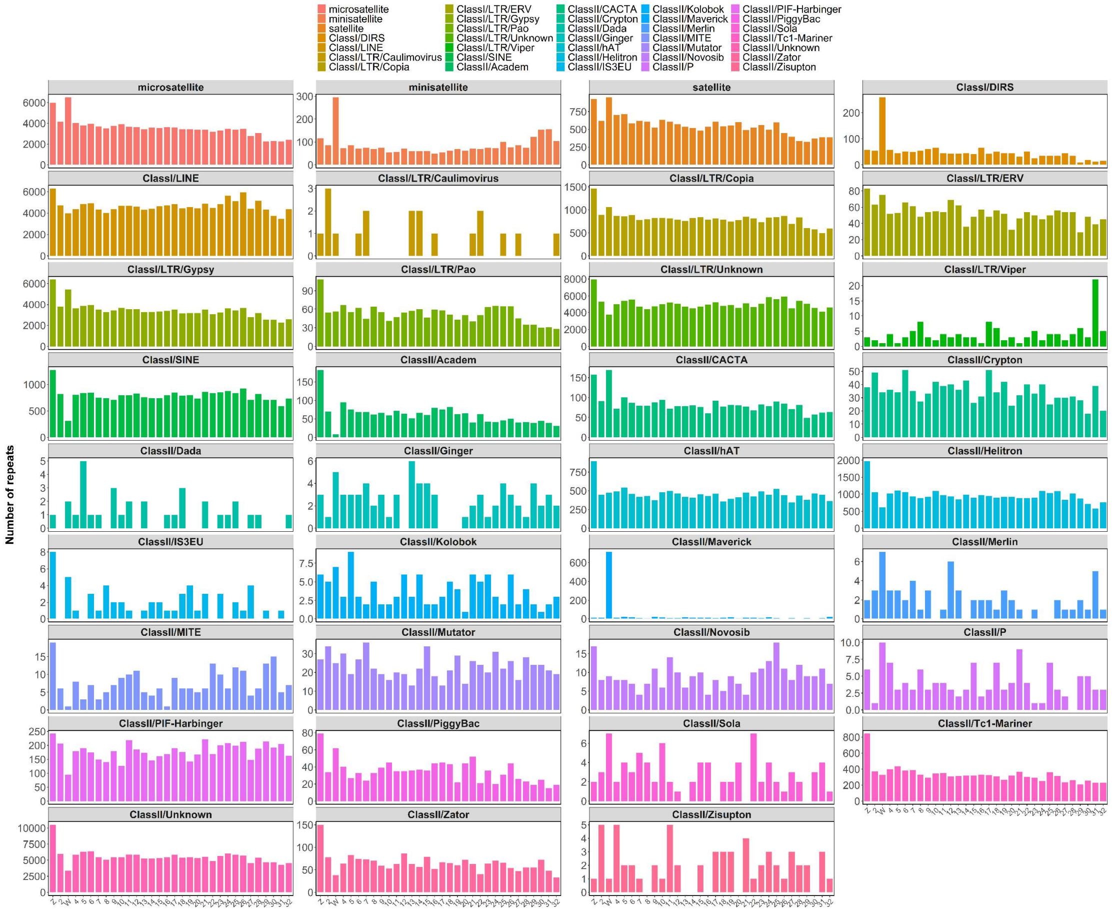


**Supplementary Figure 5. The number of repeat sequences in W chromosome (LG3).**

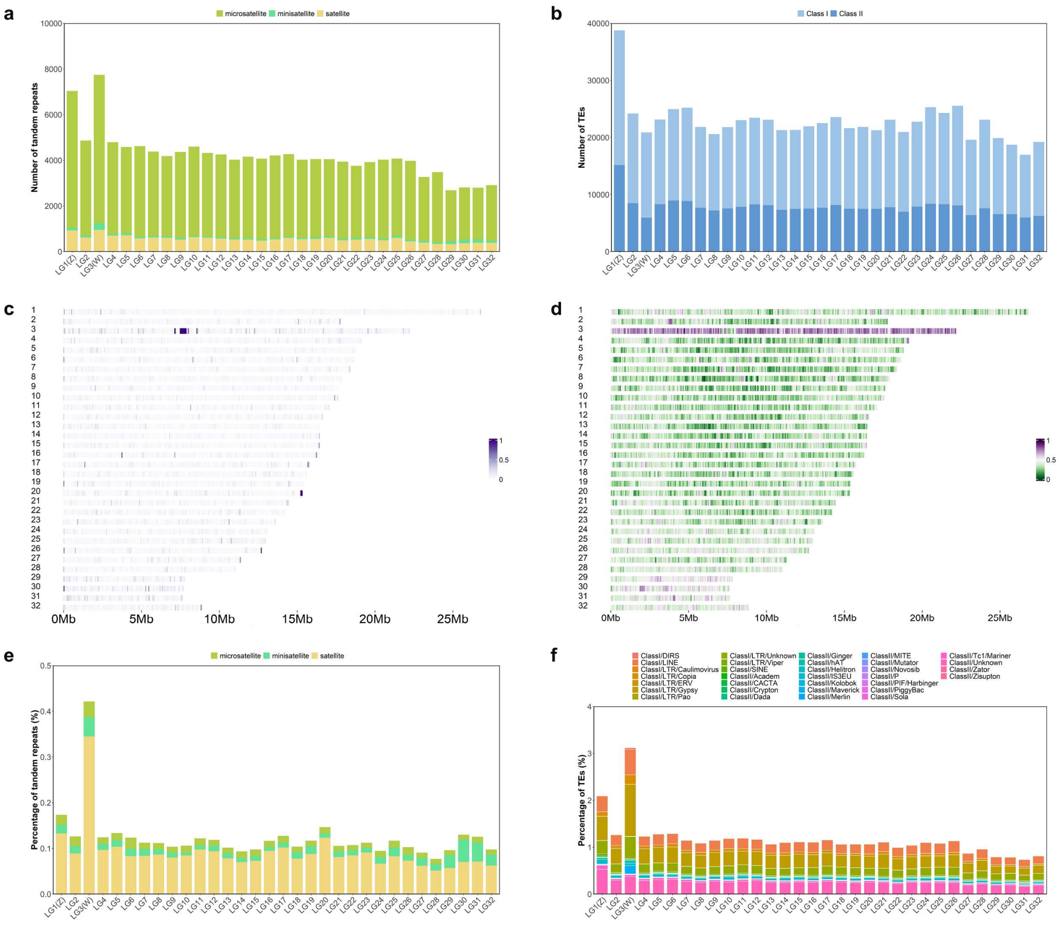


**Supplementary Figure 6. The number (a-b), density (c-d) and proportion (e-f) of repeat sequence in all chromosome (LG1-LG32).**

**Table S1. Statistics of genomic sequencing data of *Ostrinia furnacalis* by PacBio Sequel II**

| Sequence Type | Reads Num | Total Bases (bp) | Reads N50 (bp) | Mean Length (bp) | Longest Read (bp) |
|---|---|---|---|---|---|
| Subreads | 43,968,759 | 515,146,437,892 | 12,514 | 11,716 | 509,876 |
| CCS | 2,535,838 | 32,962,737,921 | 13,153 | 12,999 | 50,267 |

**Table S2. Statistics of genomic sequencing data of *Ostrinia furnacalis* by Hi-C**

Clean data by Hi-C

| ReadSum | BaseSum | GC(%) | N[a](%) | Q20[b](%) | Q30[c](%) |
|---|---|---|---|---|---|
| 177382279 | 53050949420 | 37.4 | 0 | 97.55 | 93.16 |

Mapping rate of Hi-C clean data

| Total Read Pairs | Mapped Reads | Unique Mapped Read Pairs |
|---|---|---|
| 177,382,279 | 276,188,309 | 118,446,847 |

a: The ratio of N in the bases

b: The percentage of bases with Phred quality score ≥ 20

c: The percentage of bases with Phred quality score ≥ 30

**Table S3. Statistics of genomic resequencing data of female and male pupae and transcriptome sequencing of female gonads and a mixed sample**

| Sample | ReadSum (Mb) | BaseSum (Gb) | Q30(%) |
|---|---|---|---|
| **Genomic resequencing** | | | |
| female1 | 82.2 | 12.3 | 93 |
| female2 | 94.0 | 14.1 | 94 |
| female3 | 91.4 | 13.7 | 94 |
| female4 | 92.3 | 13.8 | 93 |
| female5 | 93.0 | 13.9 | 93 |
| male1 | 93.8 | 14.0 | 93 |
| male2 | 84.4 | 12.6 | 93 |
| male3 | 95.0 | 14.2 | 93 |
| male4 | 93.7 | 14.0 | 92 |
| male5 | 87.7 | 13.1 | 93 |
| **Transcriptome sequencing** | | | |
| female_gonad1 | 21.5 | 6.4 | 94 |
| female_gonad2 | 21.2 | 6.3 | 95 |
| female_gonad3 | 20.1 | 6.0 | 95 |
| female_gonad4 | 18.1 | 5.4 | 95 |
| female_gonad5 | 19.8 | 5.9 | 96 |
| mixed sample | 42.7 | 12.7 | 95 |

**Table S4 Chromosome-level assembled Lepidoptera genomes**

| | *O. furnacalis* | *C. medinalis* | *C. suppressalis* | *C. pomonella* | *T. ni* | *H. melpomene* | *B. mori* |
|---|---|---|---|---|---|---|---|
| Genome size (Mb) | 493.1 | 528.3 | 824.4 | 772.9 | 368.2 | 269.0 | 460.3 |
| Karyotype | 2n=62 | 2n=60 | 2n=58 | 2n=56 | 2n=54 | 2n=42 | 2n=56 |
| Number of contigs | 57 | 4671 | – | 2221 | 26605 | – | - |
| Number of scaffolds | 43 | 3248 | 27144 | 1717 | 6181 | 3807 | 696 |
| Number of assembled chromosomes | 30A+Z+W | 29A+Z+W | 28A + Z | 27A+Z+W | 26A+Z+W | 20A+Z | 27A+Z |
| **Genome assembly quality** | | | | | | | |
| Contig N50 (Mb) | 15.7 | 0.5 | 0.3 | 0.9 | 0.6 | 0.1 | 12.2 |
| Scaffold N50 (Mb) | 16.5 | 16.1 | 1.8 | 8.9 | 14.2 | 0.3 | 16.8 |
| Percentage of scaffolds in chromosomes (%) | 86.8 | 89.1 | 92.5 | 97.5 | 90.6 | 82.7 | 87.3 |
| BUSCO genes (%) | 97.7 | 96.4 | 97.5 | 98.5 | 97.8 | 97.4 | 97.7 |
| **Genomic features** | | | | | | | |
| Repeat (%) | 41.5 | 39.5 | 46.4 | 42.9 | 20.5 | 24.9 | 46.8 |
| G+C (%) | 37.6 | 38.5 | 36.9 | 37.4 | 35.6 | – | 38.2 |
| **Gene annotation** | | | | | | | |
| Number of genes | 16509 | 15045 | 15653 | 17184 | 14043 | 12669 | 14623 |

**Table S5. Assessments of assembled genome**

**CEGMA**

| Number of 458 CEG* present in assembly | %of 458 CEGs present in assemblies | Number of 248 highly conserved CEGs present | % of 248 highly conserved CEGs present |
|---|---|---|---|
| 440 | 96.07% | 196 | 79.03% |

**BUSCOs**

| Complete BUSCOs(C) | Complete and single-copy BUSCOs(S) | Complete and duplicated BUSCOs(D) | Fragmented BUSCOs(F) | Missing BUSCOs(M) | Total Lineage BUSCOs |
|---|---|---|---|---|---|
| 932(97.69%) | 926(97.06%) | 6(0.63%) | 6(0.63%) | 16(1.68%) | 954 |

**DNA-seq data mapping**

| Total_reads | Mapped reads | Mapped (%) | Properly mapped (%) | Properly mapped (%) |
|---|---|---|---|---|
| 183,973,226 | 182,267,035 | 99.07% | 175,334,598 | 95.30% |

**Table S6. Genomic annotation of *Ostrinia furnacalis*.**

| | Annotated Number | Annotated Ratio |
|---|---|---|
| tRNA | 7,710 | – |
| rRNA | 73 | – |
| miRNA | 39 | – |
| pseudogenes | 167 | – |
| Annotation of protein-coding genes | | |
| GO_Annotation | 12013 | 72.77 |
| KEGG_Annotation | 12033 | 72.89 |
| KOG_Annotation | 9314 | 56.42 |
| Swissprot_Annotation | 10864 | 65.81 |
| TrEMBL_Annotation | 15901 | 96.32 |
| eggNOG_Annotation | 11746 | 71.15 |
| nr_Annotation | 16201 | 98.13 |
| All_Annotated | 16213 | 98.21 |

**Table S7. Copy number for W and autosomal/Z chromosome paralogs.**

| Aurosomes & Z | | | W chr |
|---|---|---|---|
| Gene name | Copy number | Chromosomes | Copy number |
| *algA* | 12 | LG9/LG11/LG13/LG19/LG22/LG24/LG27/LG29/LG31/LG32 | 3 |
| *Arc1* | 5 | LG16/LG21/LG25/LG26/LG28 | 3 |
| *Arc2* | 22 | LG1/LG5/LG6/LG11/LG12/LG14/LG15/LG18/LG19/LG20/LG21/LG23/LG26 | 1 |
| *At3g55350* | 8 | LG8/LG11/LG18/LG21/LG22/LG25 | 1 |
| *BZIP53* | 1 | LG19 | 1 |
| *CFDP2* | 19 | LG2/LG6/LG7/LG8/LG10/LG11/LG13/LG14/LG16/LG19/LG20/LG22/LG25/LG26/LG30/LG32 | 6 |
| *clz9* | 5 | LG1/LG8/LG11/LG26/LG32 | 3 |
| *exo* | 1 | LG21 | 3 |
| *F52C9.6* | 47 | LG2/LG4/LG5/LG6/LG7/LG8/LG8/LG9/LG10/LG11/LG12/LG13/LG14/LG15 | 4 |
| *gag* | 93 | LG1/LG2/LG4/LG5/LG6/LG7/LG8 | 9 |
| *GIP* | 15 | LG2/LG4/LG9/LG10/LG14/LG15/LG18/LG19/LG25/LG26/LG27 | 8 |
| *GIS2* | 8 | LG9/LG10/LG11/LG21/LG24/LG27/LG28 | 1 |
| *GLE1* | 4 | LG7/LG18/LG20/LG22 | 1 |
| *harbi1* | 75 | LG1/LG2/LG4/LG6/LG7/LG9/LG10/LG11/LG12/LG13 | 9 |
| *K02A2.6* | 58 | LG1/LG4/LG5/LG6/LG8/LG9/LG9/LG10/LG11/LG12/LG13/LG14/LG15 | 20 |
| *kif15-b* | 8 | LG4/LG14/LG19/LG20/LG22/LG23/LG28/LG32 | 3 |
| *L1RE1* | 14 | LG2/LG8/LG9/LG9/LG10/LG13/LG14/LG15/LG19/LG25/LG26/LG28 | 6 |
| *L1TD1* | 13 | LG2/LG5/LG6/LG16/LG17/LG18//LG19/LG24/LG28/LG32 | 20 |
| *LdOrf-130* | 17 | LG9/LG11/LG13/LG14/LG16/LG18/LG20/LG21/LG22/LG26/LG27/LG28 | 11 |
| *MSANTD3* | 54 | LG1/LG2/LG4/LG6/LG7/LG8/LG9/LG10/LG11/LG13/LG14/LG15 | 4 |
| *mutL* | 4 | LG7/LG15/LG17 | 2 |
| *NOF* | 4 | LG2/LG8/LG12/LG13 | 1 |
| *nusB* | 3 | LG11/LG14 | 1 |
| *ORF1* | 19 | LG1/LG2/LG4/LG6/LG10/LG11/LG12/LG13/LG14/LG17/LG19/LG20/LG23/LG24/LG27 | 10 |
| *panB* | 3 | LG16/LG24/LG25 | 1 |
| *PGBD3* | 11 | LG1/LG4/LG6/LG8/LG11/LG12/LG12/LG13/LG21/LG30 | 5 |
| *PGBD4* | 14 | LG1/LG4/LG10/LG12/LG13/LG16/LG20/LG21/LG22/LG23/LG25 | 3 |
| *pif1* | 9 | LG1/LG9/LG10/LG17/LG18/LG23/LG24/LG31 | 5 |

| | | | |
|---|---|---|---|
| *POL* | 113 | LG1/LG2/LG4/LG5/LG6/LG7/LG8 | 44 |
| *Pygo1* | 4 | LG4/LG5/LG22/LG25 | 1 |
| *Pym* | 2 | LG12/LG15 | 2 |
| *RAI1* | 2 | LG14/LG22 | 1 |
| *RP146* | 1 | LG25 | 1 |
| *rpe* | 4 | LG10/LG14/LG16/LG20 | 1 |
| *rpsH* | 5 | LG2/LG9/LG18/LG31 | 1 |
| *RTase* | 27 | LG2/LG5/LG6/LG8/LG9/LG10/LG11/LG12/LG14/LG16/LG17/LG18/LG19/LG21/LG22/LG24/LG28/LG29/LG31 | 7 |
| *T* | 11 | LG1/LG2/LG7/LG8/LG18/LG21/LG31 | 3 |
| *Tf2-8* | 15 | LG1/LG4/LG5/LG6/LG11/LG15/LG17/LG20/LG21/LG22/LG23/LG24/LG27 | 1 |
| *Tf2-9* | 19 | LG4/LG5/LG8/LG9/LG10/LG12/LG14/LG15/LG16/LG20/LG21/LG22/LG26/LG28/LG29/LG30/LG31 | 24 |
| *TY3B-G* | 17 | LG1/LG8/LG9/LG10/LG11/LG14/LG16/LG17/LG20/LG23/LG23/LG24/LG25/LG26 | 4 |
| *TY3B-I* | 42 | LG2/LG4/LG6/LG7/LG8/LG9/LG10/LG13/LG15/LG16/LG17/LG18 | 10 |
| *TYROBP* | 1 | LG11 | 1 |
| *Unc13c* | 25 | LG1/LG5/LG6/LG8/LG11/LG14/LG15/LG16/LG17/LG18/LG19/LG21/LG22/LG24/LG26/LG30 | 17 |
| *USO1* | 1 | LG18 | 2 |
| *valS* | 9 | LG4/LG8/LG10/LG16/LG17/LG19/LG20/LG22/LG29 | 1 |
| *ZBED1* | 4 | LG6/LG13/LG22/LG25 | 5 |
| *ZBED4* | 4 | LG21/LG25/LG31 | 3 |
| *ZC3H10* | 4 | LG1/LG11/LG26 | 2 |
